# Direct measurements of collagen fiber recruitment in the posterior pole of the eye

**DOI:** 10.1101/2023.05.07.539784

**Authors:** Po-Yi Lee, Gosia Fryc, John Gnalian, Yi Hua, Susannah Waxman, Fuqiang Zhong, Bin Yang, Ian A Sigal

## Abstract

Collagen is the main load-bearing component of the peripapillary sclera (PPS) and lamina cribrosa (LC) in the eye. Whilst it has been shown that uncrimping and recruitment of the PPS and LC collagen fibers underlies the macro-scale nonlinear stiffening of both tissues with increased intraocular pressure (IOP), the uncrimping and recruitment as a function of local stretch have not been directly measured. This knowledge is crucial for the development of constitutive models associating micro and macro scales. In this project we measured local stretch-induced collagen fiber bundle uncrimping and recruitment curves of the PPS and LC. Thin coronal samples of PPS and LC of sheep eyes were mounted and stretched biaxially quasi-statically using a custom system. At each step, we imaged the PPS and LC with instant polarized light microscopy and quantified pixel-level (1.5 μ m/pixel) collagen fiber orientations. We used digital image correlation to measure the local stretch and quantified collagen crimp by the circular standard deviation of fiber orientations, or *waviness*. Local stretch-recruitment curves of PPS and LC approximated sigmoid functions. PPS recruited more fibers than the LC at the low levels of stretch. At 10% stretch the curves crossed with 75% bundles recruited. The PPS had higher uncrimping rate and waviness remaining after recruitment than the LC: 0.9° vs. 0.6° and 3.1° vs. 2.7°. Altogether our findings support describing fiber recruitment of both PPS and LC with sigmoid curves, with the PPS recruiting faster and at lower stretch than the LC, consistent with a stiffer tissue.

**Statement of Significance:** Peripapillary sclera (PPS) and lamina cribrosa (LC) collagen recruitment behaviors are central to the nonlinear mechanical behavior of the posterior pole of the eye. How PPS and LC collagen fibers recruit under stretch is crucial to develop constitutive models of the tissues but remains unclear. We used image-based stretch testing to characterize PPS and LC collagen fiber bundle recruitment under local stretch. We found that fiber-level stretch-recruitment curves of PPS and LC approximated sigmoid functions. PPS recruited more fibers at a low stretch, but at 10% bundle stretch the two curves crossed with 75% bundles recruited. We also found that PPS and LC fibers had different uncrimping rates and non-zero waviness’s when recruited.

## 1. Introduction

Collagen is the main load-bearing component of the sclera and lamina cribrosa (LC) in the posterior pole of the eye (**Figure 1**). [1–5] Collagen thus plays a critical role in how these tissues protect the delicate neural tissues of the retina and optic nerve and maintain the globe shape necessary for vision. [6, 7] The collagen fibers of both sclera and LC exhibit the process known as fiber recruitment, [8, 9] giving rise to their well-known macro-scale nonlinear behavior (**Figure 2**). [10–14] While fiber recruitment as a function of intraocular pressure (IOP) has been measured, [14] no studies have directly measured the uncrimping and recruitment of the collagen fibers of the LC and adjacent peripapillary sclera (PPS) as a function of stretch. This knowledge is crucial to develop constitutive models of the tissues associating the micro and macro scales, which must be constructed based on the tissue stretch and not IOP. [15, 16] The knowledge is also necessary to understand the role of collagen fiber characteristics in the physiology and pathophysiology of the region. Because of the complex architecture and mechanical interactions between the PPS and LC, it is not trivial to infer recruitment as a function of stretch from recruitment as a function of IOP.

**Figure 1.**
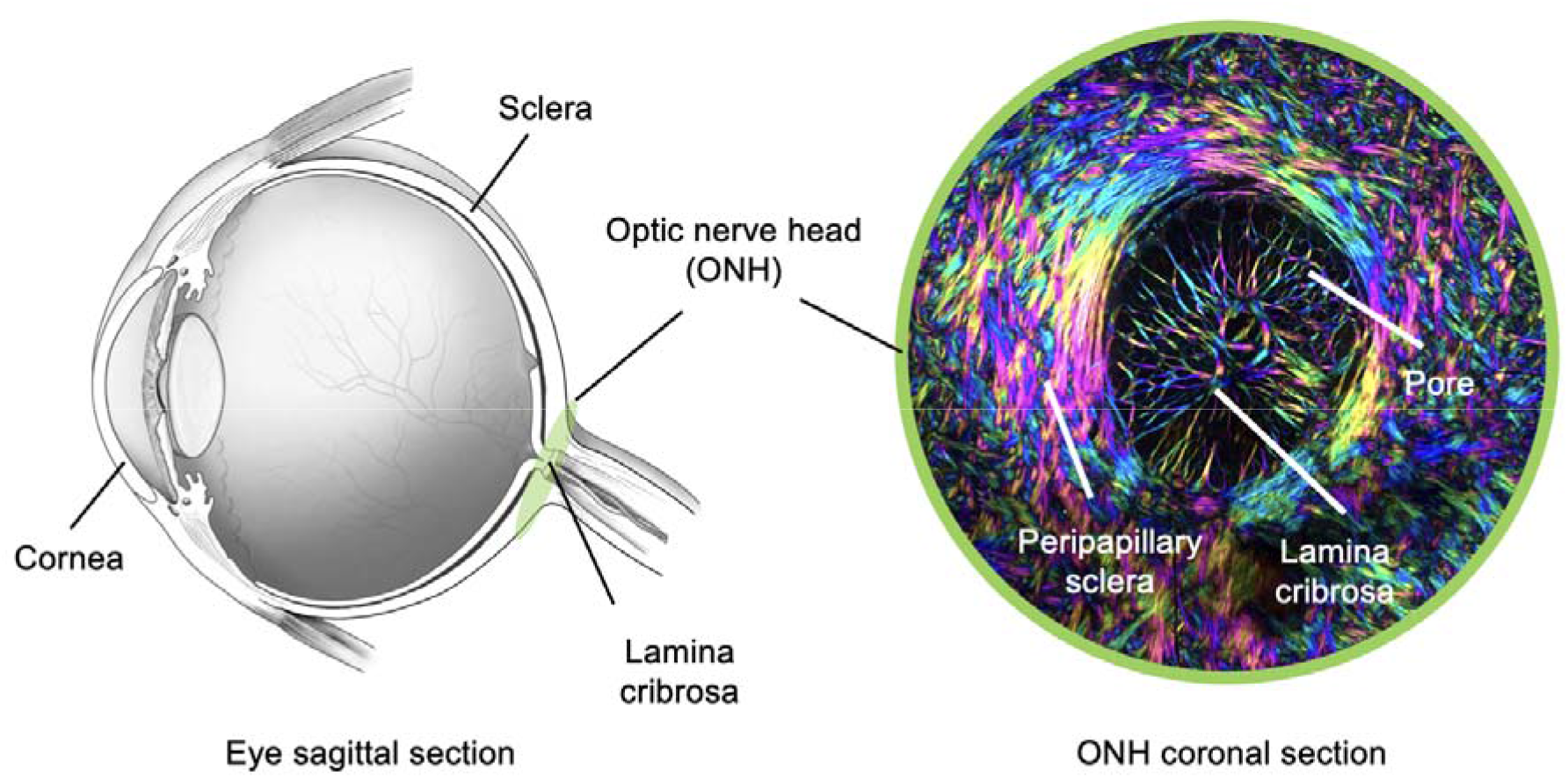
The anatomy of the eye and optic nerve head (ONH). (Left) A sagittal view of an eye is shown. Adapted from a diagram by the National Eye Institute. (Right) A coronal view of of ONH aquired by instand polarized light microscopy. The image encodes pixel-level collagen fiber orientation and density as hue and brightness, respectively. The lamina cribrosa (LC) is a mesh-like connective tissue spannign the sleral canal. Through the pores of the LC pass the retinal ganglion cell axons that carry visual information from the eye to the brain and blood vessels that perfuse the posterior pole..

**Figure 2.**
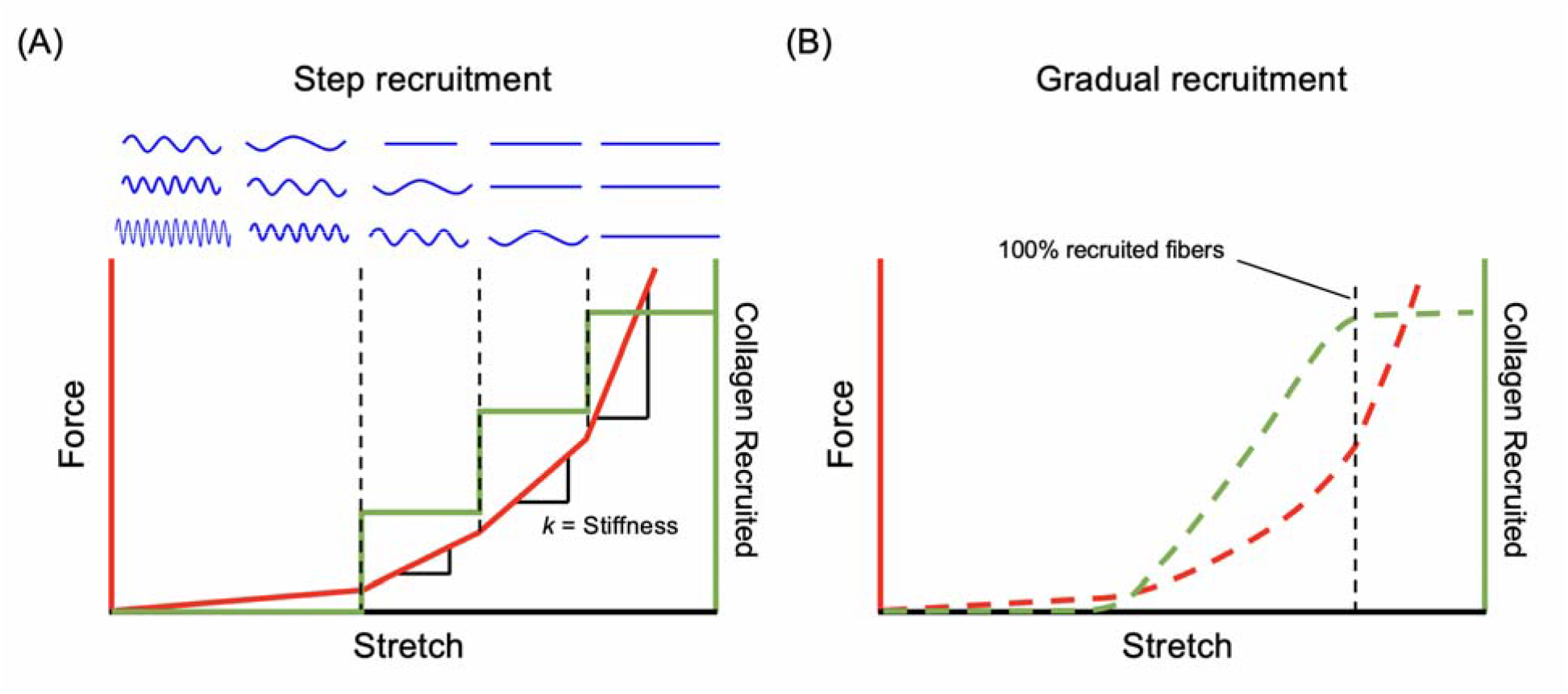
Schematic illustration of how the collagen crimp determines the nonlinear mechancial behavior of a fiber bundle. A wavy fiber is not recruited. Conversely, a fiber that is straightened is recruited. (A) In a simple recruitment model, a collagen fiber bundle consists of several fibers with variable crimp (blue curves). When no fiber is recruited, the fiber bundle has a low stretch-force slope, i.e., low stiffness. Fibers become recruited one by one as stretch increases (green curve). Step recruitment results in an abrupt, step, increase in stiffness (red curve). (B) When a fiber bundle consists of many fibers with variable crimp. A progressively increasing fraction of fibers becomes recruited as stretch increases (green curve). Gradual recruitment leads to a nonlinear increase in stiffness (red curve). When all fibers are recruited (black dash), the stiffness, i.e., stretch-force slope, becomes a constant. Actual tissues exhibit an even more complex process, with fibers rotating, interacting with one another, for instance through cross-links, are interlocked and can slide. Nevertheless, several tissues have been show to follow closely the recruitment process described here. [42, 81]

Our goal in this study was to characterize the stretch-induced collagen fiber uncrimping and recruitment of the LC and adjacent PPS at the fiber level. We hypothesized that collagen recruitment in the PPS occurs at lower levels of stretch than in the LC. This suspicion is based on three elements: First, the LC is generally thought to be more compliant than the sclera (studies suggest about an order of magnitude), [17–21] consistent with its fibers recruiting at higher levels of stretch in the LC than in the PPS. Second, experiments have shown that even at elevated IOPs the PPS suffers only small deformations (single-digit levels of strain), [2, 22] suggesting again that its fibers recruit at low levels of stretch. Third, in our measurements of LC and PPS uncrimping as a function of IOP, we found that the fibers of the sclera recruited at lower levels of IOP than those of the LC. [14] However, by using whole globes and maintaining tissue integrity, the PPS and LC interacted, allowing distortions of one tissue to affect the other. [23, 24] Because the sclera is stiffer than the LC and there is so much more sclera than LC we suspect that this likely resulted in the two tissue recruitment curves as a function of IOP being more similar than if the tissues had been tested independently. By evaluating the recruitment as a function of local stretch instead of IOP, in this study, we were able to measure directly stretch-induced fiber responses of PPS and LC and remove any potential effects from interactions with other tissues. To achieve our goals, we used a recently introduced technique, instant polarized light microscopy (IPOL), which allows measuring directly and with high accuracy the stretch-induced uncrimping at the fiber-scale without the need for labels or stains.

## 2. Methods

### A note on terminology

Ocular collagen has a complex hierarchical organization. [25–30] In some regions, such as the cornea, the hierarchy is relatively well characterized, with collagen organized into evenly-spaced micro-fibrils approximately 35nm in diameter that join to form fibrils, then fibers, then lamellae. [29, 31–33] Collagen in the PPS and LC is more complex and not nearly as well understood. The analysis in this work is based on measuring collagen features at the micro-scale revealed by polarized light microscopy such as those in Figures 4 and 5. It is at this scale that the undulations and crimp we refer to have been reported and measured. [14, 27] While the images show what look like fibers, like those in the cornea, we expect these to have a sub-structure. To acknowledge this important fact, we decided to refer to them as “fiber bundles”. In the hierarchical structure it is likely that these fiber bundles group into larger bundles.

**Figure 3.**
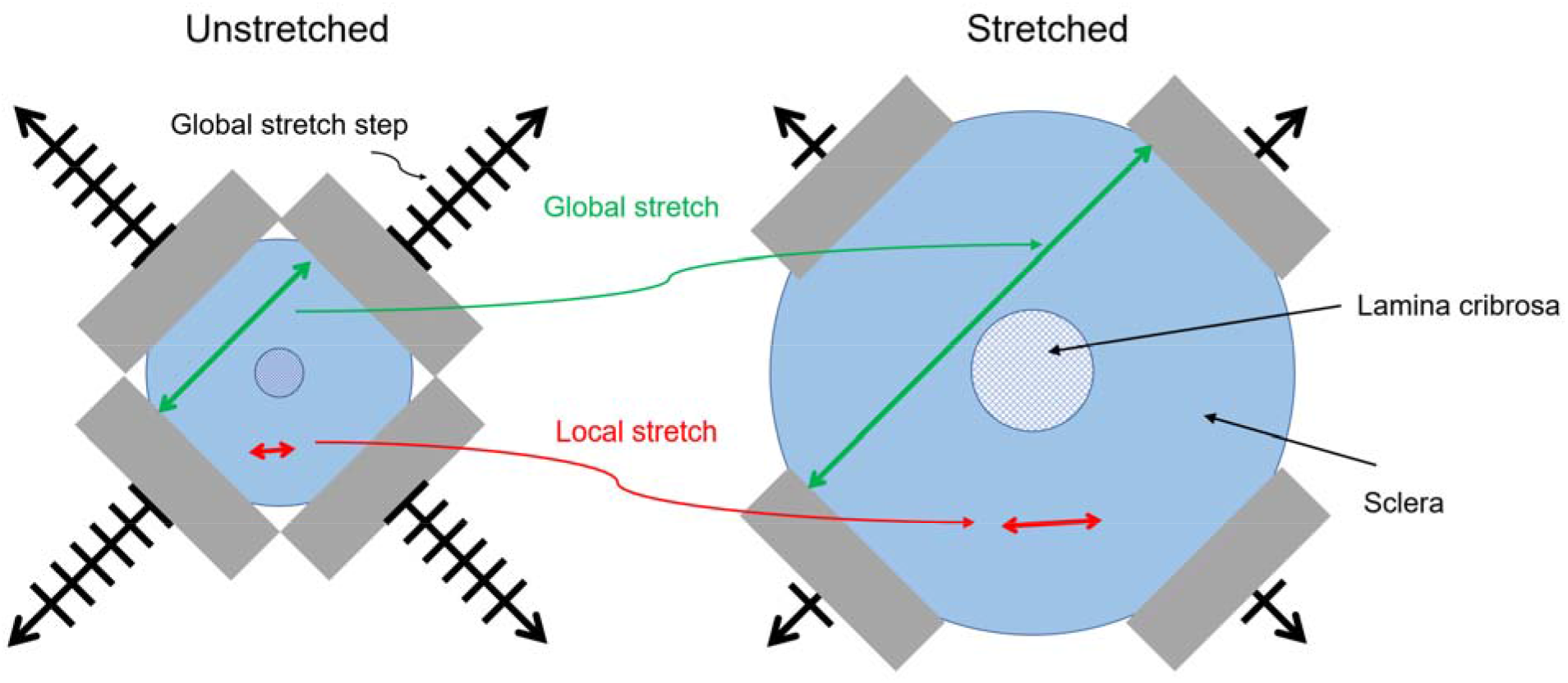
Schematic of biaxial stretch testing. Clamps (grey rentangles) mounted and stretched an ONH section (blue donut) equally along two orthogonal axes. Global stretch was defined as the change in clamp-to-clamp distance (green arrow) as the sample was stretched. Each global stretch step was 0.5 mm. Local stretch was calculated by the percent length change in the region of interest (red arrow) on the PPS or LC fiber bundles tracking by digital image correlation. Note that the amount of stretch is exaggerated for clarity.

**Figure 4.**
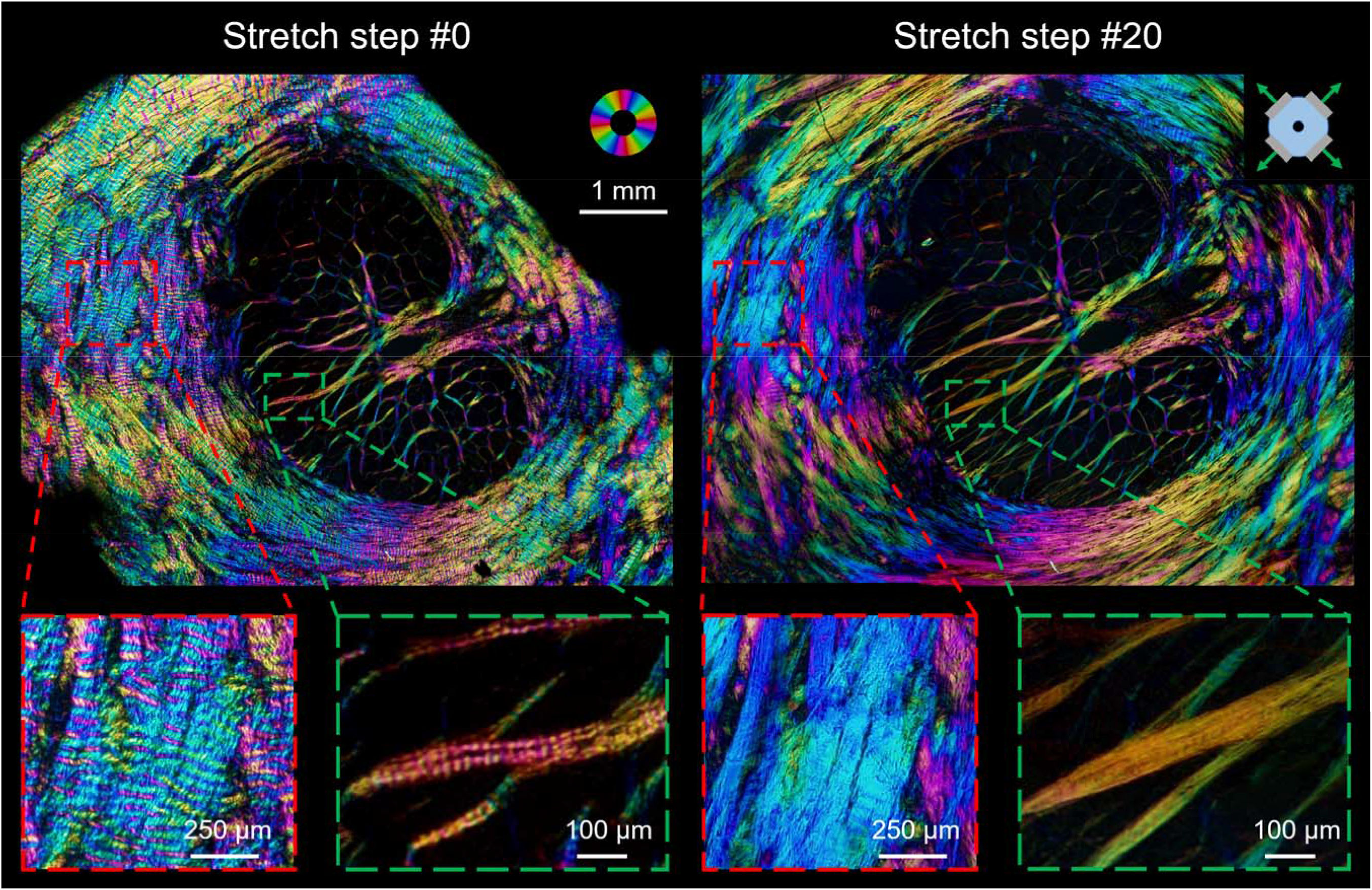
Collagen crimp visualization in the ONH. IPOL image mosaics of a coronal section from the ONH of an unfixed sheep eye under a quasi-static biaxial stretch. On the left hand as mounted before stretch (Stretch step 0). On the right hand side after stretch (Stretch step #20). The color disc on the top right-hand side of the image represents local fiber orientation, and the brightness of each pixel in the images represents local fiber density. The bottom row shows close-ups of PPS (left) and LC (right) in each condition. The dark shadows on the corners of the top left panel indicate the locations of the clamps. Using the IPOL technique allowed identifying the difference in collagen microstructure such as crimp before and after stretch. At the initial state, PPS and LC collagen fibers had clear waviness, discernible by the color bands. In contrast, at the stretched state, there were far fewer undulations in color on the PPS and LC collagen fibers, indicative of straightened, recruited fibers. A perfectly straight fiber has a single orientation and therefore a single color. Note the large changes in the canal shape, starting as elliptical and finishing close to circular.

**Figure 5.**
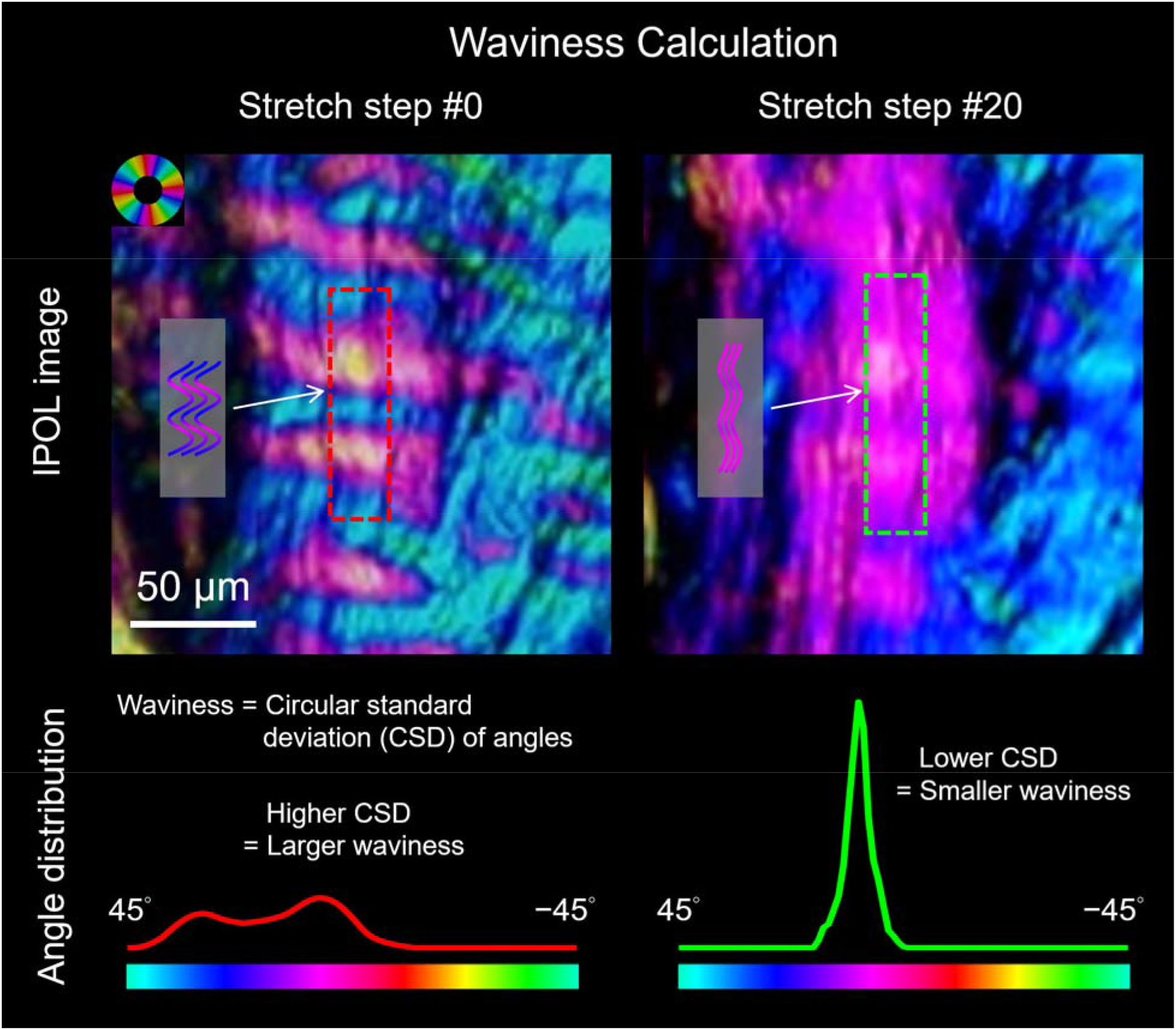
IPOL images of PPS collagen and their corresponding orientation distributions within a region of interest (dashed rectangle) at different stretch states. Waviness was calculated using the circular standard deviation of the collagen fiber orientations. In wavy crimped fiber bundles, the orientation distribution is more variable, resulting in larger waviness values (left) whereas in recruited fibers, the distribution is less variable, resulting in smaller waviness values (right). Local stretch was the percent change in the long axis between two regions of interest (red dash rectangle to green dash rectangle) on the fiber bundles.

### Sample Preparation

Three eyes from one-year-old sheep were obtained from a local abattoir and processed within four hours after death as described elsewhere. [34] Briefly, the muscles, fat, and episcleral tissues were carefully removed. The optic nerve head (ONH) region was isolated using an 11-mm-diameter trephine and embedded in optimum cutting temperature compound (Tissue-Plus; Fisher Healthcare, TX, USA). Samples were then snap frozen in liquid nitrogen-cooled isopentane and sectioned coronally at a thickness of 16 µm. The samples were then washed with PBS to remove cutting compound. The tissues were never fixed or labeled. To ensure robust analysis we made sure to have three good sections for each eye (no breaks, folds or other artifacts), *i.e.*, 9 sections from 3 sheep eyes were used and analyzed in total.

### Biaxial Stretch Testing

Each tissue section was mounted on a custom biaxial micro-stretcher system following previously reported methodology. [34] Briefly, two pieces of silicone sheeting (Medical Grade, 0.005”; BioPlexus, AZ, USA) were used to sandwich the tissue section to prevent curling or tears at the clamp points and to keep the tissue section fully hydrated. Only at the clamps was the silicone in tight contact with the tissue. The section was stretched equally along two orthogonal axes, quasi-statically followed by long (20s or more) pauses to allow dissipating viscoelastic effects in 0.5 mm stretch steps. We refer to the clamp-to-clamp stretch as global stretch (**Figure 3**).

### Imaging

Each section was imaged using IPOL following previously reported methodology to visualize the collagen. [35] IPOL allows label-free visualization and quantification of collagen structure and orientation via color information for each pixel. Briefly, a set of polarization encoder and decoder were retrofitted into a brightfield microscope, where one consisted of a polarizer and a clockwise polarization rotator and the other consisted of an analyzer and a counter-clockwise polarization rotator. A region with low or no birefringence appeared dark since the polarizations of the white light had no change and thus was blocked. A birefringent sample, such as collagen, appeared colorful since IPOL differentiated the polarizations with wavelengths and thus changed the spectrum of the white light. The colorful light was acquired by a color camera to produce the true-color images indicating collagen fiber orientation. [36]

IPOL was implemented with a commercial inverted microscope (IX83; Olympus, Tokyo, Japan), a color camera (DP74, Olympus, Tokyo), and a 4x strain-free objective (numerical aperture 0.13, 1.5 μm/pixel). At each stretch step, multiple images were captured to cover the entire ONH region in a mosaic. Images with 20% overlap were acquired using a translational stage and stitched into mosaics using Fiji. [37] We have shown that the visualization of collagen fibers is not affected by mosaics or stitching. [27, 38] Autofocus was applied to avoid out-of-focus images during the stretch.

### Collagen Crimp Quantification

IPOL is based on the optical properties of the collagen molecules, and therefore it allows accurate quantification of collagen orientation without the need to resolve and distinguish fibrils, fibers, or bundles. Pixel-scale collagen fiber orientations were determined by matching hue values over a pre-calibrated color-angle conversion map for corresponding angles. [34] In IPOL images, color stripes along the ONH fiber bundles indicate crimp (**Figure 4** **left**); as the stretch increased, disappearing stripes indicate decreased crimp (**Figure 4** **right**). In this work, we measured the distribution of orientations along collagen crimp, and used these to determine biomechanical behavior through the stretch-induced changes in crimp.

A custom MATLAB program was developed for manually placing rotatable rectangular regions on PPS and LC along crimp directions. As a measure of crimp, we computed “waviness” defined as the circular standard deviation of the pixel-level fiber angles within the rectangular region. For wavy crimped fiber bundles, the orientation distribution was bimodal and wide, indicated by higher waviness values (**Figure 5** **left**). For straight, or recruited, fiber bundles, the distribution was unimodal and narrow, indicated by smaller waviness values (**Figure 5** **right**). We tracked the long axis of the rectangular region using digital image correlation and then updated the region based on the elongation and displacement of the long axis from the tracking result at each step [39]. We then measured the waviness in the updated rectangular region at each global stretch step until tissue failure. Local stretch was calculated by the length change in the long axis of the rectangular region (**Figure 3**). From the waviness-stretch data, we determined if a fiber bundle was recruited when the waviness no longer decreased with stretch, and then measured “% stretch to recruit” and “waviness when recruited” of recruited fiber bundles. We then used linear regression on the waviness-stretch data before recruitment to obtain the “uncrimping rate” as the absolute magnitude of the slope *k* (**Figure 6**). The rationale and implications of the definition of fiber recruitment and the waviness when recruited measurement are addressed in the Discussion. Note that we observed that some PPS and LC fiber bundles did not recruit with the stretch. We suspect that this was in cases where tissue processing had broken the fibers, disconnecting them from one another and/or the matrix. In the local stretch analysis, we focused on the collagen fiber bundles that did recruit.

**Figure 6.**
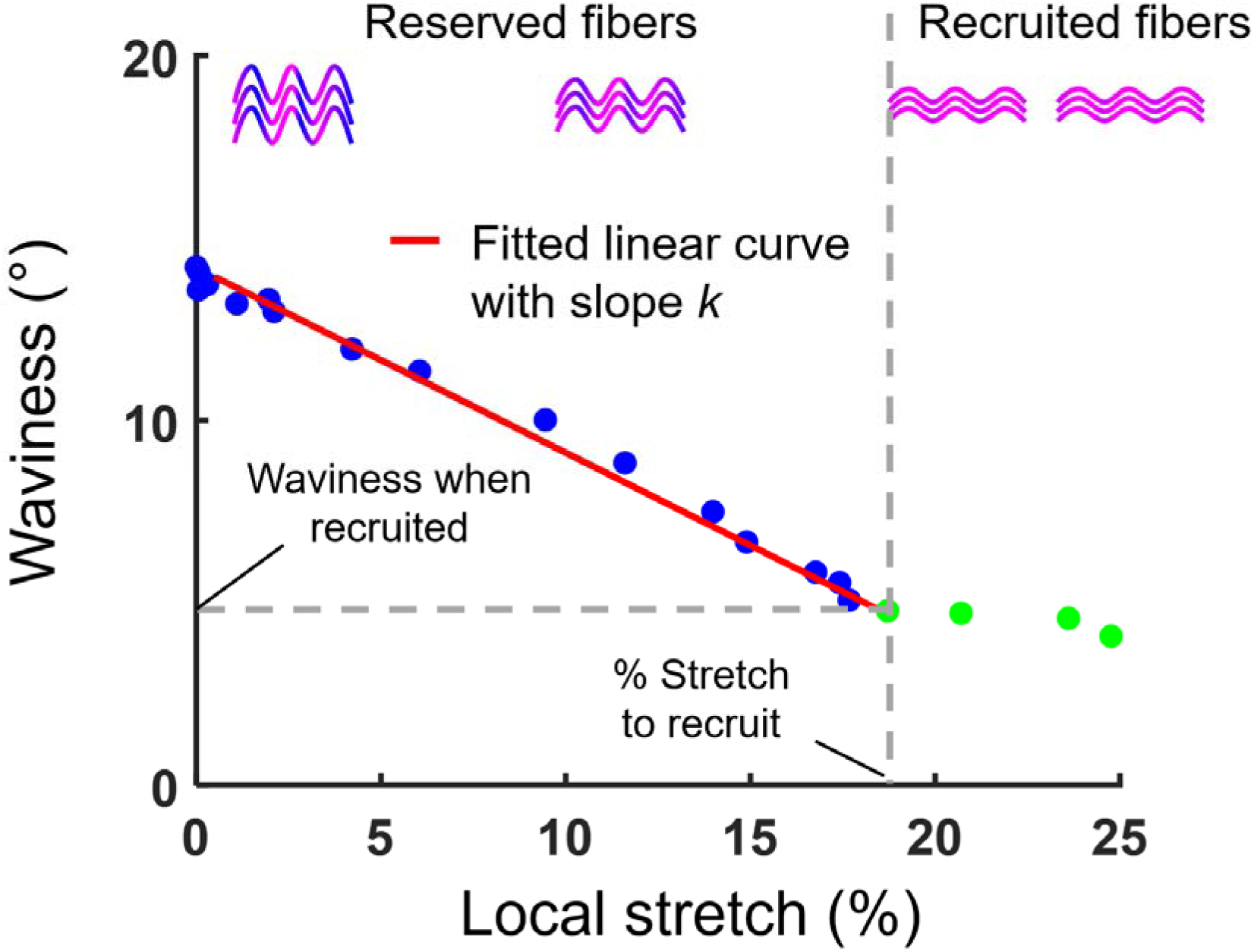
Local waviness-stretch curve. Local stretch represents the elongation in the local region. This is different from the clamp-to-clamp biaxial stretch of the entire ONH. As the local stretch increased, the waviness of collagen fibers of the posterior pole decreased. Fiber recruitment was defined as when the waviness of collagen fibers remains constant despite further increases in local stretch. The values at that point are the “waviness when recruited” and “% stretch to recruit”. The waviness-stretch curve until recruited was fitted using linear regression. The absolute magnitude of the slope *k* of the fitted line was defined as the “uncrimping rate”.

### Statistical analysis

Linear mixed effect models accounting for correlations between measurements from the same sections and eyes were used to analyze the differences in waviness between the PPS and LC at each stretch step, and the differences in waviness when recruited and uncrimping rate between the PPS and LC. Descriptive statistical calculations such as angular mean and standard deviation were made using circular statistics. [4, 40] All statistical analyses were done using R. [41]

## 3. Results

We obtained the percentages of fiber bundles recruited at given local stretch in the LC and PPS (**Figure 7a**). In the recruited fiber bundles, we found that the uncrimping rate in the PPS (0.9±0.5) was significantly higher than that in the LC (0.6±0.3) (*P*’s < 0.001) (**Figure 7b**). We found that the PPS recruited waviness when recruited (3.1°±0.9°) was significantly higher the LC waviness when recruited (2.7°±0.7°) (*P*’s < 0.05) (**Figure 7c**).

**Figure 7.**
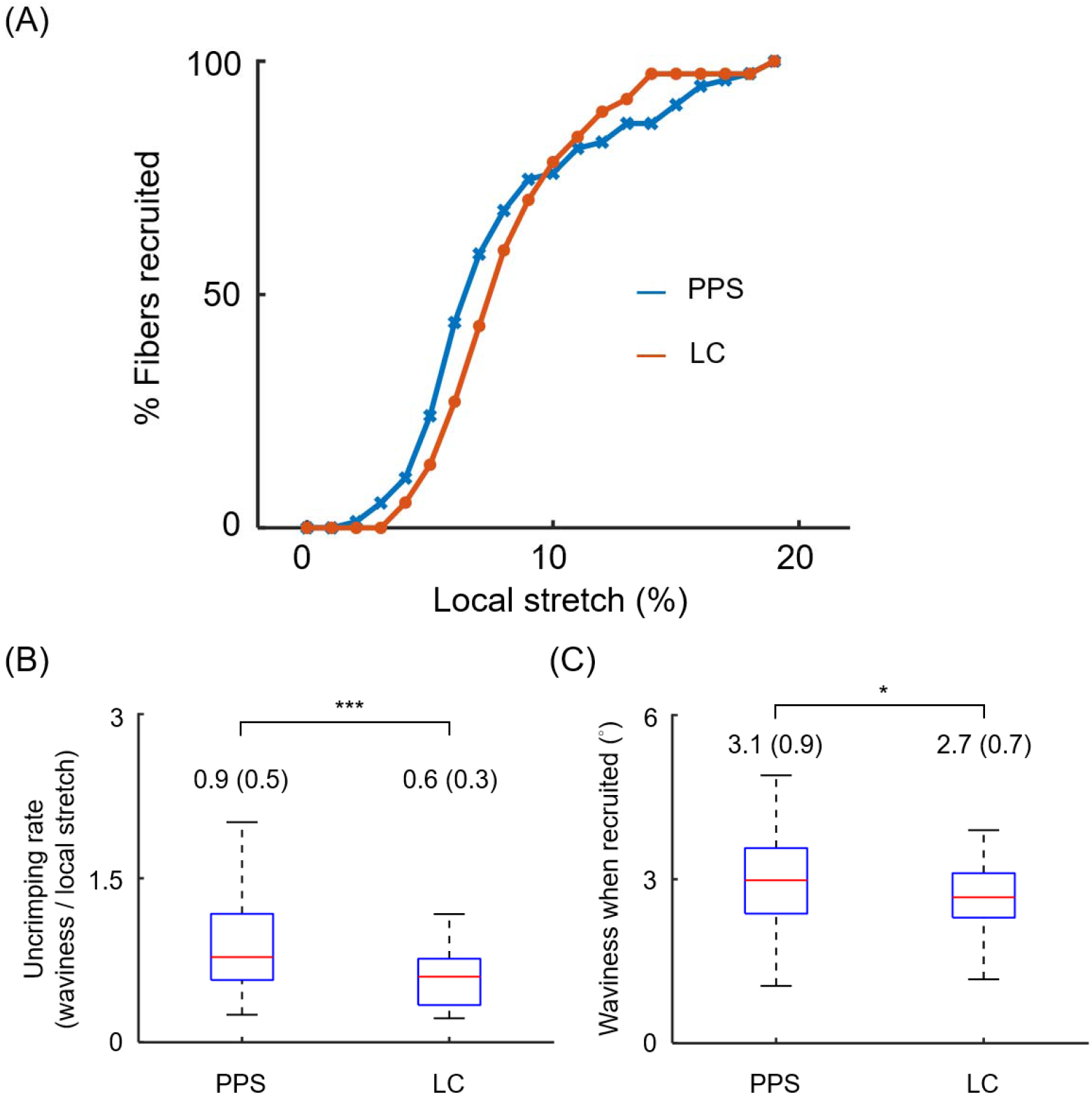
Collagen fiber recruitment curves of PPS and LC under local stretch. (A) A total of 75 PPS collagen bundles and 37 LC beams were recruited under stretch testing. Local stretch-recruitment curves are plotted based on the measurements “% stretch to recruit” of PPS and LC. Both curves approximated a sigmoid function. About 75% of PPS and LC fibers were recruited at a 10% local stretch. The PPS recruited more fibers than the LC when the local stretch was less than 10%. (B) Boxplot of uncrimping rates in the PPS and LC beams. Average (standard deviation) uncrimping rates for PPS and LC beams are labeled above the whisker’s tip. PPS beams had a significantly higher uncrimping rate (*P*’s < 0.001). (C) Boxplot of waviness when recruited in the PPS and LC beams. Average (standard deviation) waviness when recruited for PPS and LC beams are labeled above the whisker’s tip. The waviness when recruited of the collagen fiber in the PPS was larger and more variable than that of the LC (*P*’s < 0.05).

Waviness of both PPS and LC decreased with global stretch (**Figure 8**). At the initial state, the waviness was significantly higher in the PPS (10.4°±2.7°) than in the LC (8.3°±2.6°) (*P*’s < 0.001). The difference was no longer significant between PPS (3.5°±1.2°) and LC (3.3°±1.2°) after the tissues were stretched (*P*’s < 0.001).

**Figure 8.**
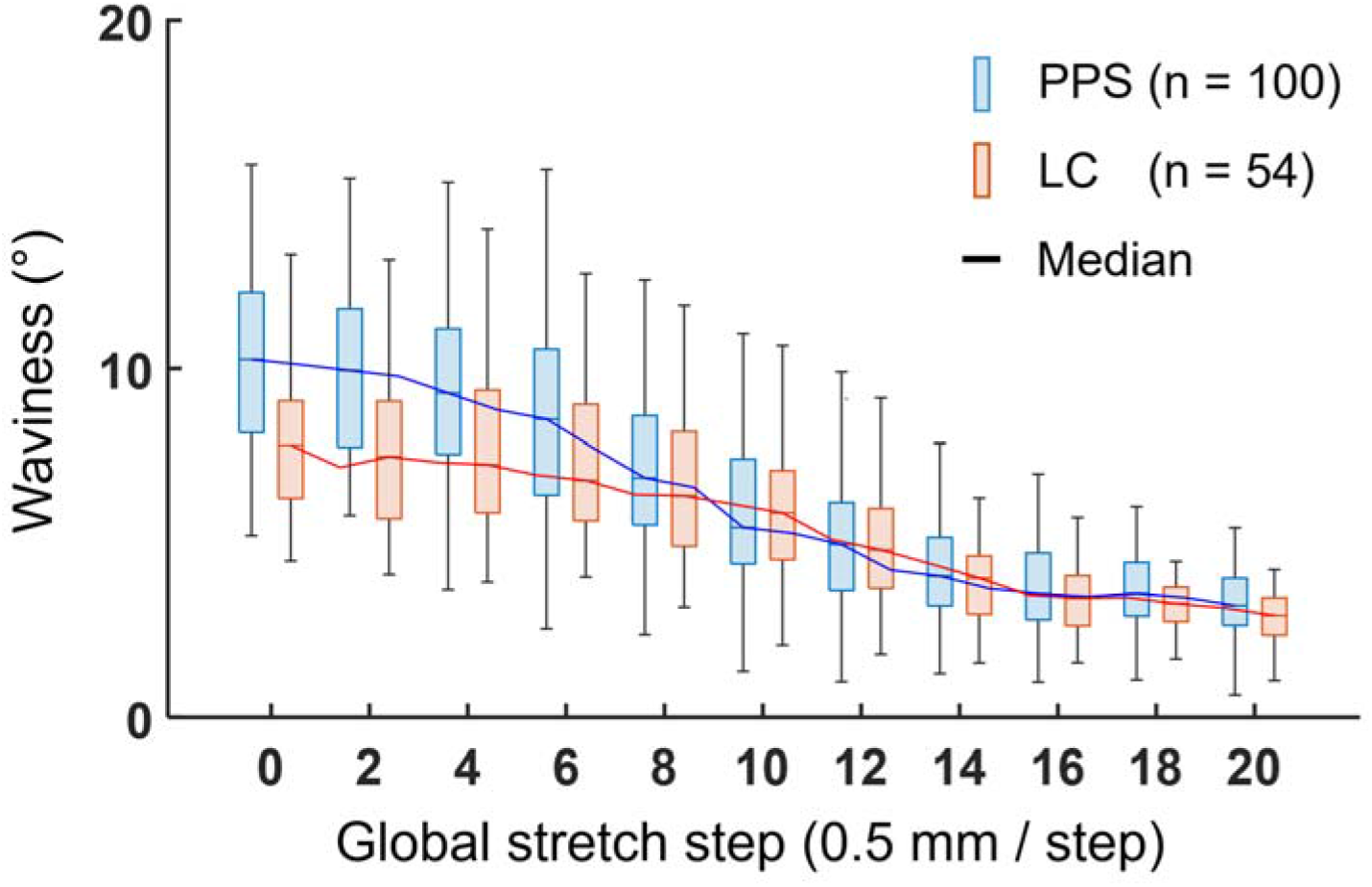
Collagen waviness of the PPS and LC decreased with global stretch. Each stretch step was 0.5 mm for an ONH section isolated using an 11-mm-diameter trephine. At each stretch step, we measured the distributions of the collagen crimp waviness in the PPS (blue) and LC (orange), represented here as thin box plots. The median of the waviness at each stretch step were connected to highlight the differences between PPS and LC. A total of 154 measurements of waviness were collected, of which 100 were of PPS collagen bundles and 54 of LC beams across all sections. At the initial state the average (SD) waviness measurements were 10.4° (2.7°) for PPS beams and 8.3° (2.6°) for LC beams, showing that the waviness in the PPS was significantly higher than the waviness in the LC (*P*’s < 0.001). At the fully stretched state, the average (SD) waviness measurements were 3.5° (1.2°) for PPS beams and 3.3° (1.2°) for LC beams, showing that there was no significant difference between the waviness in the PPS and LC.

## 4. Discussion

We applied image-based stretch testing to measure local stretch-induced collagen fiber bundle uncrimping and recruitment of the PPS and LC. Three main results arise from this work. First, PPS fiber bundles recruit at lower levels of local stretch than LC fiber bundles but both tissue recruitment curves were similar. Second, PPS fiber bundles had a higher uncrimping rate than LC fiber bundles. Third, waviness’s when recruited in the PPS and LC were non-zero. Below we discuss each of these in turn.

### PPS fiber bundles recruit at lower levels of the local stretch than LC fiber bundles but both tissue recruitment curves were similar

Only recruited fibers bear substantial loads, [14, 27, 42–44] and therefore the local stretch-recruitment curves reveal key information on the relative fraction of potentially load-bearing fiber bundles that are actually bearing the load. The results in this work reveal that a larger fraction of PPS fiber bundles bear the load when the stretch is small (<10%), a difference in the rate of regional stiffening under stretch. This is consistent with the general impression that the LC is more compliant than the sclera. [21] This supports the concept that PPS stiffening at a lower level of stretch could be a protective mechanism that allows the PPS to resist large deformations at low and moderate IOPs, relieving the LC loads and protecting the fragile nerve fibers in the LC from vision-threatening insult. The phenomenon that recruitment implies that not all fibers are contributing to bear loads is well characterized in other soft tissues, like tendon and ligament. [45, 46]

While the findings support our hypothesis that collagen recruitment in the PPS occurs at lower levels of stretch than in the LC, we were surprised by the similarity between the recruitment curves for both tissues. Similar curves indicate similar rates of PPS and LC stiffening under stretch. We recently conducted a study in which we used computational modeling to estimate recruitment curves as a function of IOP in seven regions of the corneoscleral shell. [47] As expected, all regions exhibited sigmoid curves, but the recruitment rates varied substantially over the globe. We found that the fibers of the posterior equator recruited the fastest at low IOPs, such that at 15mmHg over 90% of the fibers were recruited. The fibers of the PPS recruited much slower with IOP, such that only 34% of the PPS fibers were recruited at a physiologic IOP of 15 mmHg. At an elevated IOP of 50mmHg, 70% of the PPS fibers had been recruited. Experimental studies of IOP-induced sclera deformation have shown that even at elevated IOPs, the PPS undergoes single digit levels of stretch. [11, 48, 49] Together, those results are consistent with this work in that we measured 75% recruitment in PPS at 10% local stretch.

The recruitment curves for PPS and LC cross at 10% stretch, with both tissues exhibiting 75% recruitment. It is interesting to compare this finding with those in a recent study in which we analyzed crimp in the LC and PPS. [14] The study was similar to this one, but with two important differences: we measured crimp as a function of IOP, not as a function of stretch as in this work. The study was cross-sectional, using samples fixed at specific IOPs. Despite the differences, the results from both studies are remarkably consistent with one another. In the previous study, the recruitment curves of the PPS and LC were similar, as in this work, with the PPS exhibiting higher recruitment at IOPs under 15mmHg. Interestingly, both curves crossed at a physiologic IOP of 15mmHg, with 75% of collagen fibers recruited in both PPS and LC. The recruitment curves are slightly different at higher IOPs or higher levels of stretch, although both studies coincide in that there was substantial collagen waviness in both the PPS and LC even at very high IOPs.

An important consideration when interpreting our findings is that the recruitment curve is only related to the rate of stretch-induced stiffening, which may be very different from the actual tissue stiffness. The results above should not be interpreted as indicative of which tissue is stiffer. In addition to the crimp angle, a computational modeling approach [44] has shown that the elastic modulus of the filament *E* and the ratio of the amplitude of the helix to the radius of the filament cross-section *R_0_*/*r*_0_ affect stretch-dependent stiffening of collagen. Interestingly, they found that, everything else being equal, the initial crimp angle has almost no influence on the overall shape of the stress and stiffness function. In a later study, Grytz and colleagues used computational modeling to infer micro-scale fiber crimp and mechanics from macro-scale experimental data on posterior pole inflation.[11] They found that collagen fibril strain and recruitment rate were similar between the peripapillary and the mid-peripheral regions at the micro-scale, but the in-plane strains were significantly higher in peripapillary region than in the mid-periphery.

In this study, we considered tissues as part of the LC or PPS, without further details of their location within the ONH. In an earlier cross-sectional study we analyzed the collagen crimp in eyes fixed at 0, 5 or, 10mmHg IOP. [9] Crimp was characterized by period, not waviness as in this work. Elsewhere we have discussed the relationship between these two measures [14, 27] We found small, uniform, crimp periods within the LC, and immediately adjacent PPS. The crimp period in the PPS increased with distance from the canal. We interpreted those findings as indicative that the tissues of the ONH are adapted to prevent very different deformations at the scleral canal edge region that could insult the neural tissues. Our findings in this work are consistent with those observations.

### PPS fiber bundles had a higher uncrimping rate than LC fiber bundles

Uncrimping rate is closely associated with the shape of the crimp. [50] Physically-based structural models have been developed to describe how the configuration of collagen evolves under load and predict the non-linear mechanical behavior of collagen fiber microstructures that undergo large deformations. [29, 50–54] Collagen crimp can be viewed as an extensible helical spring, i.e., a rod wound around the central axis of a cylinder with a radius *R_0_*. [44, 55] In the structural model, a plane sinusoidal crimp [*R_0_* = 0] has a higher uncrimping rate than a helical crimp [*R_0_* > 0]. [50] PPS fiber bundles having a higher uncrimping rate is consistent with the collagen configuration of PPS being closer to a plane sinusoidal structure. In geometry, a plane sinusoidal structure allows for “packing” more collagen in a unit volume than a helical structure. This is supported by IPOL images (**Figure 4**) that the density of collagen in the PPS regions is higher since PPS regions in IPOL images are brighter than LC regions. This is also consistent with the general impression that the LC is more compliant than the PPS. [21] It is worth noting that PPS fiber bundles had a higher uncrimping rate but the “% stretch to recruit” of PPS was lower at higher stretch. It is because the waviness of the collagen fibers in the PPS was larger and higher variable than that of the LC at the initial and early stretching states (**Figure 8**). The high uncrimping rate led to the waviness of collagen fibers in the PPS being close to that of the LC after the tissues were stretched. However, even though the uncrimping rate was higher in the PPS, some PPS fiber bundles with high waviness still required more local stretch to achieve recruitment.

### Waviness’s when recruited in the PPS and LC were non-zero

Waviness when recruited relates to the ability of a fiber bundle to bear the load and resist being fully uncrimped under stretch. Ideal fiber recruitment happens when a fiber is fully straightened (*i.e.*, zero waviness). However, interactions between fibers may prevent the fibers from being fully straightened. For example, stretch cannot fully remove fiber twisting and bending and thus such fibers always present residual crimp under large stretch (**Figure 9**). High waviness when recruited thus could indicate a more complex fiber microstructure. Moreover, it is worth noting that an elevated load could potentially increase the waviness of these microstructures. The examples show only uniaxial loading, which is known to lead to fiber alignment with the load, through rotation, if possible, bending if not. The tissues of the PPS, and to some extent those of the LC, are subjected to biaxial loading. Studies using inflation have even suggested that at the macro-scale the sclera is sometimes close to an equi-biaxial stretch. [56] At the micro-scale, the situation is more complex. This could explain the findings from our previous study, where the recruitment percentages of PPS and LC decreased for IOP increases between 20 mmHg and 50LmmHg, as the recruitment was determined based on waviness thresholds in that study. [14] The such microstructure has been recognized in fibrillar-level tendon and ligament, which fulfill local fibril flexibility and behave as biological hinges both absorbing tension and guiding the recoil of collagen fibers. [57, 58] The interactions between fibers that result from the complex microstructure and interweaving can also have major effects at the tissue macro-scale. [3]

**Figure 9.**
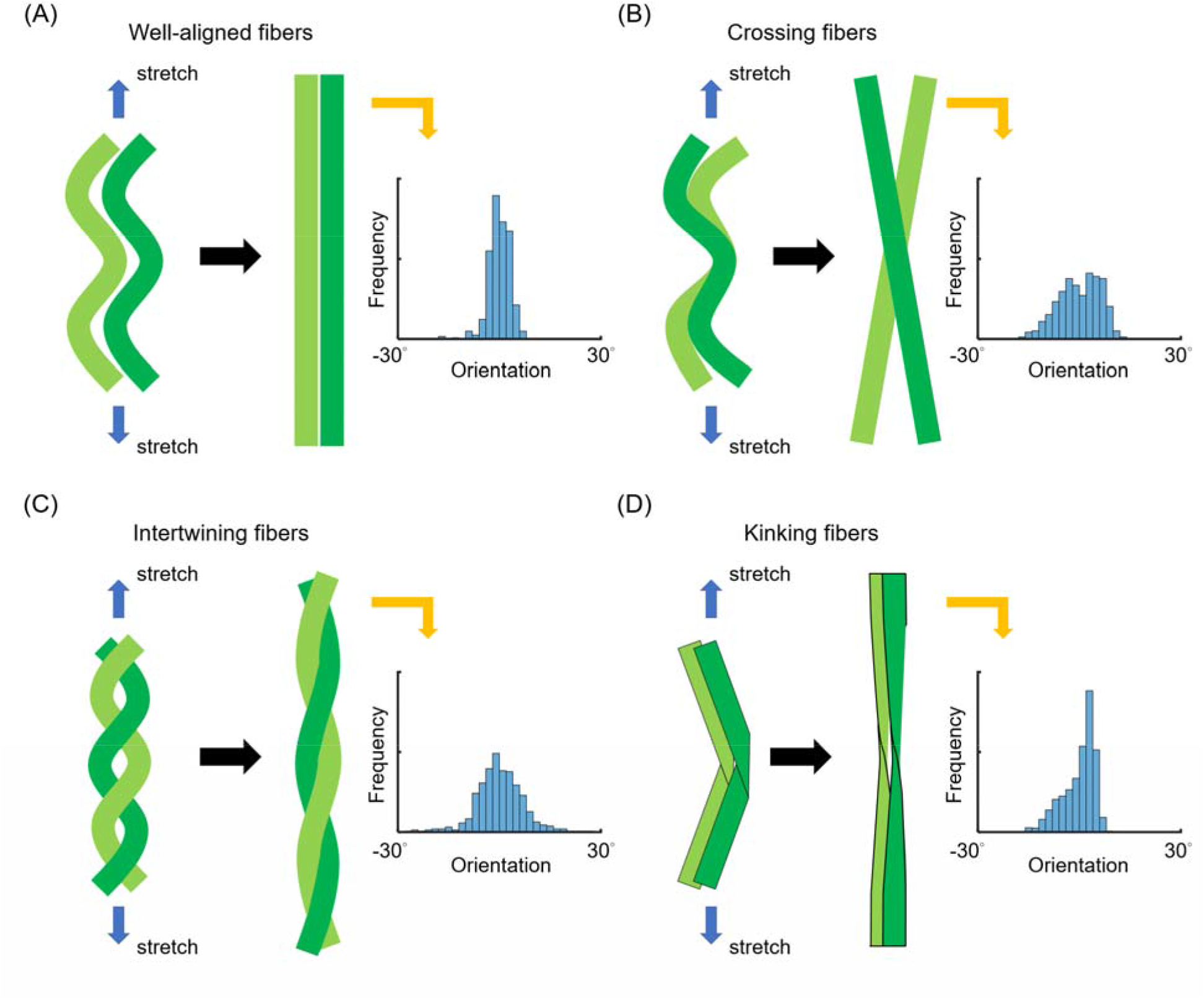
Illustrative examples of how the microstructure at the fiber scale can result in non-zero waviness even under full recruitment. (A) Ideal fiber recruitment indicates that fibers are well-aligned, with a narrow distribution of orientations and a near-zero waviness. When fibers are (B) crossed (i.e., a distribution with two peaks), (C) intertwined (i.e., a wide distribution), or (D) kinked (i.e., a skewed distribution), fibers never lose their structural identity under stretching and thus still present residual waviness even when fully loaded and maximally recruited. In such cases, waviness when recruited is larger than zero. Therefore, waviness when recruited and its corresponding angle distribution may reflect how a fiber microstructure differs from a well-aligned one. Note that despite the excellent resolution of IPOL, it is still not sufficient to directly identify such sub-microstructures.

It is important to note that the relationship between waviness and local stretch measurements may be different from the relationship between waviness and global or macro-level stretch. Global stretch often results in varying local stretch, leading to also varying different levels of recruitment and stiffening between regions. Local stretch measurements allow tracking the continuous uncrimping process of an individual fiber bundle. These measurements exclude macro-level sample displacements and deformations during testing, such as unfolding, rotations, and slip.

In a pervious cross-sectional study, we evaluated collagen fiber recruitment based on analyzing crimp differences using different samples fixed at specific IOPs. [14] We had to rely on analyzing the whole set of places and the baseline similarity between samples. With the techniques of the previous study, we would only be able to measure the statistical behavior of the whole set. There are some aspects of the behavior that we could not detect in the previous study but can detect now, such as the uncrimping rate and waviness when recruited of individual fiber bundles.

To the best of our knowledge, we report herein the first systematic direct measurements of collagen fiber uncrimping and recruitment in the PPS and LC while under biaxial stretch. Conventional methods to characterize collagen uncrimping processes such as cross-sectional studies [14] are considered indirect since they are not based on continuous tracking and measurement of the same real sample under increased loads. Similarly, inverse models, [44] while powerful, are also not direct measurements. Direct measurements allow identifying collagen crimp morphology when the individual fiber bundles recruit and the relationship between collagen crimp morphology and stretch. Such information will help develop more accurate constitutive models to describe ocular tissue mechanics. [59–61] Direct measurements of stretch-induced change in crimp help understand the nonlinear mechanical behavior of the posterior pole and building the connection between tissue stiffness and aging/age-related diseases in the eye. [62–64]

In this study, we used a polarized light imaging technique, introduced in 2021, IPOL, to visualize collagen fiber orientation and microstructure via a color snapshot. [35] IPOL has high spatial and angular resolution, making it uniquely well-suited to discern challenging aspects of the ONH collagen microarchitecture, such as crimp (**Figure 5**). This is particularly important for LC and PPS that have relatively small crimp, that cannot be measured well using tracing techniques that are common in vascular tissues. [42] Current versions of polarization sensitive optical coherence tomography [65] or wide-angle X-ray scattering [66] do not have sufficient spatial or angular resolution. MRI can discern PPS crimp, [67] but the complexity and cost of the system and the need to use a so-called magic angle technique makes it impractical at this moment. Some studies have used imaging techniques with higher magnifications such as confocal and multiphoton imaging [68, 69], but these have much smaller fields of view, complicating analysis. Further, they often require substantial tissue processing. A crucial strength of IPOL for this work is that it does not require labels or stains, making the process simpler and avoiding steps that could affect tissue structure and/or mechanics. IPOL also has a high temporal resolution, limited only by the frame rate of the camera. Although we had to use IPOL with mosaicking to capture a wide region, the frame rate of IPOL was still faster than polarized light imaging techniques that require multiple polarization acquisition [38] and scanning-based imaging techniques such as second harmonic imaging. [70] For this particular work faster imaging was helpful to prevent tissue dehydration during stretch testing

Readers should I consider the limitations of our study and our analysis in this work. First, we studied sheep eyes, which are similar to but not identical to human eyes. [71]Sheep eyes have a collagenous LC, but differ, among other things, in having a ventral groove in the ONH, similar to that in pig. [72] Though it is possible that the stretch-induced ONH collagen deformation found in sheep is not the same in humans, it is nevertheless still important to understand sheep as an animal model. [73, 74] Future work should include additional animal models as well as human eyes. The sheep from which we obtained the eyes were young and healthy, as is to be expected from an abattoir. We have shown that crimp in the eye decreases with age, [4] and thus it is likely that older eyes behave differently.

Second, while the architecture of collagen fibers in the eye is three-dimensional, we have limited the sample to a two-dimensional section. Stretching tissue sections is a notable simplification that likely affects stretch-induced collagen deformations. It is crucial to acknowledge that this is not a condition intended to represent the physiological state. It is extremely difficult to figure out the mechanisms of collagen recruitment in a 3D sample due to complicated structural and mechanical interactions in 3D. A possible approach is to use inflation to apply realistic loads and deformations to the fibers, but without a suitable imaging technique the tissues have to be fixed and thus crimp can only be studied at one IOP. This is the approach we followed before. Direct measurement of fiber uncrimping under stretch is critical to ensure that the mechanisms of uncrimping and stiffening are accurate. [29, 75, 76] We contend that our setup is useful for micro-mechanical testing to understand tissue fundamental mechanics. By isolating the tissue section, we were able to visualize and analyze quantitatively in-plane fiber bundles. The signals of collagen fibers decrease with fiber inclination (*i.e.*, out-of-plane orientation). [77] Therefore, analyzing the bright regions allowed identifying the fibers primarily aligned in the section plane as well as the stretching plane. In this sense, we were able to visualize the fibers that are responding to the load.

Third, biaxial stretch testing is an approximation to simulate the effects of IOP or other loads. The global stretch is different from physiological loading. Although inflation experiments can be applied to whole eye globes and are more physiologic, currently it is not possible to track the uncrimping processes due to the limited imaging resolution and penetration. Techniques leveraging structured illumination may allow overcoming these limitations, [78] but they are still in development. Overall, our methods should be thought of as providing measurements that are realistic for the local micro-mechanical fiber behavior, such as local stretch-induced uncrimping, and not of the macro-scale or global behavior. This distinction is a fairly common assumption in studies on ONH biomechanics when the intent is to simulate the tension in the ONH induced by IOP. [60, 79, 80] It remains to be determined whether the ONH collagen deformation identified in this study extends to physiological loading conditions or not. Also, whilst it is fairly likely that the loading applied to the scleral samples is not physiologic, the conditions may be more realistic for LC beams. Because the neural tissues adjacent to the beams are substantially more compliant than the beam, the assumptions of in-plane stretch along the beam may approximate the physiological condition better than in PPS.

Lastly, an important goal of this work was to provide direct measurements of collagen fiber recruitment to help drive fiber-based constitutive model formulations for the posterior pole. Although we obtained local stretch-recruitment curves, we did not measure the stresses and strains within the PPS and LC. Our results will have to be integrated with other studies to derive the constitutive models.

### In conclusion

We quantified the changes in collagen waviness in the PPS and LC under local stretch. We derived tissue-specific recruitment curves and reported several parameters including waviness when recruited and uncrimping rate, which encode information on tissue microstructure and crimp types. We found that while the PPS fibers recruit at lower levels of stretch than the LC, the two tissues have similar recruitment curves. Both curves cross at about 10% stretch with 75% collagen fiber recruitment. Interestingly, in a previous cross-sectional study of recruitment as a function of IOP we had found that curves for PPS and LC cross at normal IOP (15 mmHg) and 75% recruitment. Altogether, our findings have demonstrated that image-based stretch testing allows for characterizing microstructural collagen crimp properties of the posterior pole.

## Disclosures

### Funding

Supported in part by National Institutes of Health R01-EY023966, R01-EY028662, P30-EY008098 and T32-EY017271 (Bethesda, MD), the Eye and Ear Foundation (Pittsburgh, PA), and Research to Prevent Blindness (unrestricted grant to UPMC ophthalmology, and Stein innovation award to Sigal IA).

Fuqiang Zhong was at the University of Pittsburgh when he contributed to this work. Other authors have nothing to disclose.

## Notes

### Competing Interest Statement

The authors have declared no competing interest.

